# Loss of TCF-1 regulates production of noncanonical Tregs in a prion-like manner

**DOI:** 10.1101/2021.03.11.435008

**Authors:** Rebecca Harris, Mahinbanu Mammadli, Mobin Karimi

**Author notes:** To whom correspondence should be addressed: Mobin Karimi, Assistant Professor of Immunology and Microbiology, SUNY Upstate Medical University, 766 Irving Ave Weiskotten Hall Suite 2281, Syracuse, NY 13210, Laboratory.

## Abstract

Regulatory T cells (Tregs) are suppressive immune cells used for a variety of clinical and therapeutic applications. Canonical Tregs express CD4, FOXP3, and CD25, which are considered definitive markers of Treg status when used together. However, a subset of noncanonical Tregs expressing only CD4 and FOXP3 have recently been described in some infection contexts. The transcriptional regulation of these cells is still unclear. We found that loss of TCF-1 in all T cells in mice leads to expansion of these cells in multiple tissues in a cell-intrinsic fashion. This effect was not due to aberrant expression of FOXP3, as other functional Treg markers were also expressed. In addition, presence of TCF-1-deficient cells in a chimeric mouse induced increased production of noncanonical Tregs from WT donor cells. Therefore, targeting of TCF-1 may remove suppression on this Treg lineage, increasing the yield of these cells for use in the clinic.

**Summary sentence:** Loss of TCF-1 causes expansion of CD25- FOXP3+ noncanonical Tregs, and TCF-1-deficient T cells induce increased production of CD25- Tregs from WT cells.

## Introduction

T cells are critical for production of an immune response against many types of threats. CD8+ T cells are cytotoxic cells that recognize specific peptide/MHC I complexes, whereas CD4+ T cells recognize specific peptide/MHC II complexes and provide help to developing immune cells^1^. Overactivation of T cells may lead to a state of chronic inflammation and immunopathology, so several mechanisms are in place to avoid inappropriate activation of T cells. The main mechanism that governs appropriate T cell reactivity is thymic education, which involves positive selection and negative selection of T cells^2^. This process generates T cells that react weakly to self MHC/peptide complexes (positive selection) but deletes T cells that recognize self MHC/peptide complexes too strongly (negative selection)^2, 3, 4^. However, a small fraction of T cells may avoid negative selection, which could potentially lead to autoimmunity. When this occurs, peripheral mechanisms act to prevent the activation of these potentially autoreactive T cells^2, 5^. One such mechanism involves the suppression of T cell activation by regulatory T cells (Tregs), a subset of T cells with inhibitory function^6^. The importance of Tregs in protection against autoimmunity is exemplified by the systemic fatal autoimmunity that develops in Scurfy mice^7, 8^, which are deficient in Tregs, as well as in patients with IPEX (immune dysregulation, polyendocrinopathy, and X-linked) syndrome^9^. Scurfy mice and IPEX syndrome patients harbor a mutation in FoxP3^7, 8, 9^, which is essential for Treg development and function^10^. Thus, the presence of FoxP3+ Tregs is critical for the induction of peripheral tolerance.

Canonical Tregs are identified by expression of FOXP3 along with CD25, but reports have identified a noncanonical population of Tregs which is FOXP3+ CD25-^11, 12^. Unlike developing cells in the thymus, which can also express FOXP3 but not CD25, these noncanonical Tregs are found in the periphery lacking expression of CD25. These noncanonical Tregs are suppressive in several experimental models, including autoimmune encephalitis, inflammatory bowel disease, allergy, and diabetes models^13, 14, 15, 16, 17^. Despite the observation of these cells in many context, it is still unknown exactly which transcription factors drive production of these Tregs. However, CD25-deficient mice are unable to produce suppressive Tregs, suggesting that CD25 expression is required at some point for suppressive ability of CD25- Tregs, and is later lost in these cells^13^.

Tregs help to maintain tolerance and immune homeostasis by inhibiting the proliferation of and cytokine production by other T cells^6, 10, 18^. It is uncertain exactly how Tregs suppress other cells, but a variety of mechanisms have been described. These include: cell-contact dependent suppression (such as through Treg-expressed CTLA-4 binding to CD28 on the target cell), release of inhibitory cytokines (i.e. IL-10 and TGF-b), adsorption of IL-2, or modulating dendritic cell co-stimulation^18^. Using quantitative modeling, a recent study has also suggested that the major mechanism of Treg suppression is through a reduction in division destiny – the number of total divisions before a cell becomes quiescent – rather than reduced proliferation or increased cell death^18^. Although the functional mechanisms of these cells are still unclear, Tregs are known to express CTLA-4, IL-10, and TGF-b when they are functionally capable of suppression^19, 20, 21, 22^.

T Cell Factor-1 (TCF-1) was recently shown to play a role in Treg survival when in combination with LEF-1^23^. TCF-1 has also recently been identified as a suppressive factor of FOXP3, as it binds to the promoter region of FOXP3 to prevent aberrant expression of FOXP3 in conventional T cells^24^. Delacher et al. [2020] used viral overexpression of TCF-1 in T cells to show that TCF-1 suppresses induction of FOXP3 in conventional T cells under Treg-inducing culture conditions. In addition, they showed that mice with global TCF-1 deficiency had an increased frequency of CD25- FOXP3^int^ T cells, and that FOXP3 expression was also increased among CD8 T cells. These data, combined with the finding that these cells did not express CTLA-4 as expected, led the authors to suggest that loss of TCF-1 leads to aberrant FOXP3 expression in T cells, not to expansion of true Tregs. Finally, the authors used CRISPR to knock out TCF-1 in human and mouse CD4 T cells, and found an increase in FOXP3+ CD4 T cells^24^.

However, TCF-1 is a critical T cell transcription factor for T cell development, CD4/CD8 lineage maintenance, and responses to infection^25, 26, 27, 28, 29, 30, 31, 32^, so global deficiency of this factor drastically influences T cell development and function^33^. It is unknown whether TCF-1 deficiency in mature T cells has a similar effect on FOXP3. The potential for off-target effects of CRISPR systems is also well-documented^34, 35, 36^, leading to potentially faulty conclusions on the true role of TCF-1 in controlling mature T cell expression of FOXP3. Our studies instead utilized a T cell-specific deletion of TCF-1^37^ to investigate the effects of TCF-1 loss in mature cells on canonical (CD25+ FOXP3+) and noncanonical (CD25-FOXP3+) Tregs. We found that loss of TCF-1 in mature T cells led to an increased in frequency and number of noncanonical, FOXP3+CD25- Tregs, with no impact on to frequency but a drop in numbers of canonical CD25+ Tregs. This effect was cell-intrinsic, and unaffected by changes to Eomes and T-bet, which are altered when TCF-1 is lost. Eomesodermin (Eomes) and T-box transcription factor TBX21 (T-bet) are downstream of TCF-1, and also play critical roles in maintaining T cell lineage^32, 38, 39^.

Of critical importance, when WT cells were present in the same microenvironment as TCF-1-deficient Tregs (in a chimeric mouse), TCF cKO T cells induced elevated production of noncanonical Tregs from WT donor cells. Therefore, TCF cKO T cells have a prion-like ability to promote noncanonical Treg fate among nearby cells, despite the WT phenotype of these neighboring cells. These CD25- Tregs were found in multiple tissues, and were not expanded due to aberrant expression of FOXP3, because other functional Treg markers (CTLA-4 and IL-10) were also expressed in these cells. IL-2 production was maintained in TCF cKO CD25- Tregs at similar levels to WT CD25- and CD25+ Tregs. CD8 T cells expressing FOXP3 without CD25 were not increased by loss of TCF-1. Thus, our results show that while TCF-1 does appear to control FOXP3 expression, loss of TCF-1 in mature T cells does not simply cause FOXP3 to be expressed aberrantly. Instead, TCF-1 deficiency induces expansion of a unique CD25-FOXP3+ subset of Tregs which has recently been described in multiple models. Our observation of a prion-like trait of TCF-1-deficient T cells to induce this phenotype in WT cells within the same microenvironment is also novel. These studies show that TCF-1 modulation could be used to affect noncanonical Treg production and yield, and may lead to a prion-like ability to affect the fate of other nearby cells in the microenvironment.

## Materials and Methods

### Mice

Thy1.1 (B6.PL-Thy1a/CyJ, 000406), B6-Ly5 (CD45.1+, AKA “WT” or B6.SJL-Ptprc^a^ Pepc^b^/BoyJ, 002014), and BALB/c mice (CR:028) were purchased from Charles River or Jackson Laboratory. TCF cKO mice (Tcf7 flox/flox x CD4cre)^37^ were obtained from Dr. Jyoti Misra Sen at the NIH and bred in our facilities. CD4cre (022071), Eomes flox/flox (017293), and T-bet flox/flox (022741) mice were purchased from Jackson Laboratories. CD4cre mice were bred in our facilities with Eomes or T-bet flox mice to produce Eomes cKO or T-bet cKO mice, respectively. All mice used for transplants were female, and flow cytometry experiments were done with both male and female mice. All animal experiments were approved by the IACUC at SUNY Upstate Medical University. All procedures (including animal maintenance) were performed according to the rules and guidance provided by the IACUC. Mice aged 8–12 weeks were used, and all experiments were performed with age and sex-matched mice.

### Flow cytometry

To analyze expression of Treg markers, lymphocytes were collected and stained for flow cytometry. Lymphocytes were obtained from organs, filtered with a 70uM filter, and treated with RBC Lysis Buffer to remove red blood cells (RBCs). The cells were then washed with ice-cold MACS buffer (1x PBS with EDTA and 4g/L BSA) and plated in a 96-well V-bottom plate. Antibody cocktails were prepared in 1x PBS or MACS buffer and added to each well. The cells were stained for 30 minutes on ice, covered to protect from light. The cells were then spun to remove antibodies and washed 1-2 times with ice-cold 1x PBS or MACS. Cells were fixed overnight at 4C in 200uL of fixative (Fix/Perm Concentrate and Fixation Diluent from FOXP3 Transcription Factor Staining Buffer Set, eBioscience cat. No. 00-5523-00). The next day, stained cells were permeabilized by washing twice with permeabilization buffer (eBioscience cat. No. 00-5523-00) and then stained for intracellular markers for 40min at room temperature with antibody in perm buffer, covered from light. Cells were then washed 1-2 times with perm buffer, resuspended in 200-400uL of FACS buffer (eBioscience cat. No. 00-4222-26), and transferred to flow tubes. Data was collected on a BD LSRFortessa cytometer (BD Biosciences). Data were analyzed using FlowJo v9 (Treestar).

### Antibodies

All antibodies were purchased from eBiosciences, Biolegend, or BD Biosciences. Antibodies used included: anti-CD4-BV785, anti-FOXP3-APC, anti-CD25-PE, anti-CD25-BV421, anti-CD45.2-PE/Cy7, anti-CD45.1-Pacific Blue, anti-Thy1.1-AF700, anti-Thy1.2-APC, anti-CD4-FITC, anti-CD45.1-PE, anti-CD3-APC/Cy7, anti-CD8-FITC, anti-CD8-PE, anti-IL-2-PE/Cy7, anti-CTLA-4-PE, and anti-IL-10-APC/Cy7. LIVE/DEAD Fixable Aqua Dead Cell Stain (Invitrogen cat. No. L34957) was used to remove dead cells from the analysis. Anti-CD3 (clone 17A2, Biolegend cat. no. 100202) was used to coat stimulation plates, and Ultra LEAF-purified anti-CD28 (clone 37.51, Biolegend cat. no. 102116) was used as a soluble stimulator.

### Chimera Production

To produce bone marrow chimeras, Thy1.1 female mice aged 8-12 weeks were lethally irradiated with 800 cGys in a single dose. Bone marrow was isolated from femur and tibia of WT (B6Ly5, CD45.1) and TCF cKO (CD45.2) mice, filtered through a 70uM filter, and counted. Bone marrow cells were mixed at a 1:4 (WT:TCF cKO) ratio to ensure survival of KO cells with a potential proliferation defect. Bone marrow cells were suspended in sterile 1x PBS for transplantation. The mixed bone marrow was injected into the Thy1.1 recipients via the tail vein at 4 hours post-irradiation. At 9 weeks post-transplant, blood was collected from the Thy1.1 mice and tested via flow cytometry for presence of both CD45.1 (WT) and CD45.2 (TCF cKO) cells via flow cytometry. At 10 weeks post-transplant, recipient mice were euthanized and splenocytes were obtained for flow cytometry as described above.

### Isolation of lymphocytes from liver

To isolate lymphocytes from liver, recipient mice were euthanized and the livers were perfused with 5 mL of cold 1x PBS. The livers were then mashed through a 70uM filter, washed with MACS buffer, and mixed with 10% Percoll in RPMI/PBS. The samples in Percoll were spun at 2200rpm for 22min at 22C, with no brake or acceleration, leading to isolation of lymphocytes in the pellet. These lymphocytes were then treated with RBC Lysis Buffer to remove RBCs, and processed for flow cytometry as described above.

### Isolation of lymphocytes from small intestine

To isolate lymphocytes from the small intestine, mice were euthanized, and following liver perfusion as described above, the entire small intestine was removed and placed in ice-cold media. The guts were cut open lengthwise to expose the lumen, and washed with ice-cold media. The tissue was then placed in a 50mL tube with 20mL of strip buffer (containing 1x PBS, FBS, EDTA 0.5M, and DTT 1M) and shaken at 37C for 30 min. This removes the epithelium from the guts, so following incubation, the tissues were vortexed, supernatent was discarded, and the tissue was moved to a new tube. The gut was minced into small pieces, and 10mL of digestion buffer was added to each tube (containing collagenase, DNAse, and RPMI). The tissues were incubated for 30 min, shaking at 37C, and filtered into a clean tube using a 70uM filter. Remaining tissue pieces were ground onto the filter to obtain any remaining lymphocytes while leaving fat and structural cells behind. Finally, these cells were spun down in Percoll as described for livers, to isolate lymphocytes. RBC Lysis Buffer was not used on these cells.

### Cell culture for Treg markers

To test cells for the Treg markers (FOXP3, CD25, CTLA-4, IL-2, IL-10), splenocytes were obtained from WT or TCF cKO naive mice. These cells were split into three groups. Group 1 was simply stained for flow cytometry. Group 2 was cultured in LAK media for 6 hours with Brefeldin A (GolgiPlug), in wells previously “coated” with PBS as a control. Finally, Group 3 was cultured in LAK media for 6 hours with Brefeldin A in wells coated with anti-CD3 in PBS (1ug/mL), and anti-CD28 (2ug/mL) was added to the LAK media for culture. After 6 hours of culture, all cells were collected from the stimulation plate and stained for flow cytometry. All cells were stained for extracellular markers, then fixed overnight with the Invitrogen Intracellular Fixation and Permeabilization buffer kit (cat. no. 88-8824-00). The next day, the cells were permeabilized and stained with intracellular markers (FOXP3 for all, IL-2 and IL-10 for cultured cells only). All samples were then run on a BD LSRFortessa flow cytometer as described above.

### Statistics

All statistics were performed using one-way ANOVA, two-way ANOVA, or Student’s t-test, depending on the dataset. ANOVA analyses included Tukey’s multiple comparisons test. P-values are presented as <0.05 being significant. Data graphing and statistical testing was performed with GraphPad Prism v7 or v9 (GraphPad Software, San Diego, CA). Data are presented as means with standard deviation. All experiments were done with at least 3 mice per group, according to power analyses, and repeated multiple times.

## Results

### Loss of TCF-1 in all T cells leads to increased production of noncanonical Tregs

Delacher et al. [2020]^24^ recently claimed that loss of TCF-1 induces aberrant FOXP3 expression due to release of suppressive control. However, this work was done with TCF-1 global KO mice or CRISPR-mediated deletion in T cells, giving potential off-target effects or developmental changes in these cells. This work also did not address the role of TCF-1 in mature T cells. Therefore, we sought to determine whether loss of TCF-1 specifically in mature T cells would alter Treg populations. We obtained TCF-1 cKO mice which are deficient in TCF-1 in all T cells following the DP stage of development. We phenotyped these TCF-1-deficient mice, and found that loss of TCF-1 resulted in no change to canonical, FOXP3+ CD25+ Tregs [Fig 1A-B]. However, TCF cKO mice did have a significant increase in the frequency of noncanonical, FOXP3+ CD25- Tregs [Fig 1A-B]. These cells were not increased in CD4cre control mice [Fig 1A], so the expansion of these cells was due to loss of TCF-1, not due to the CD4cre system for deletion. Therefore, TCF-1 normally suppresses noncanonical Tregs, and loss of TCF-1 relieves this suppression.

**Fig 1.**
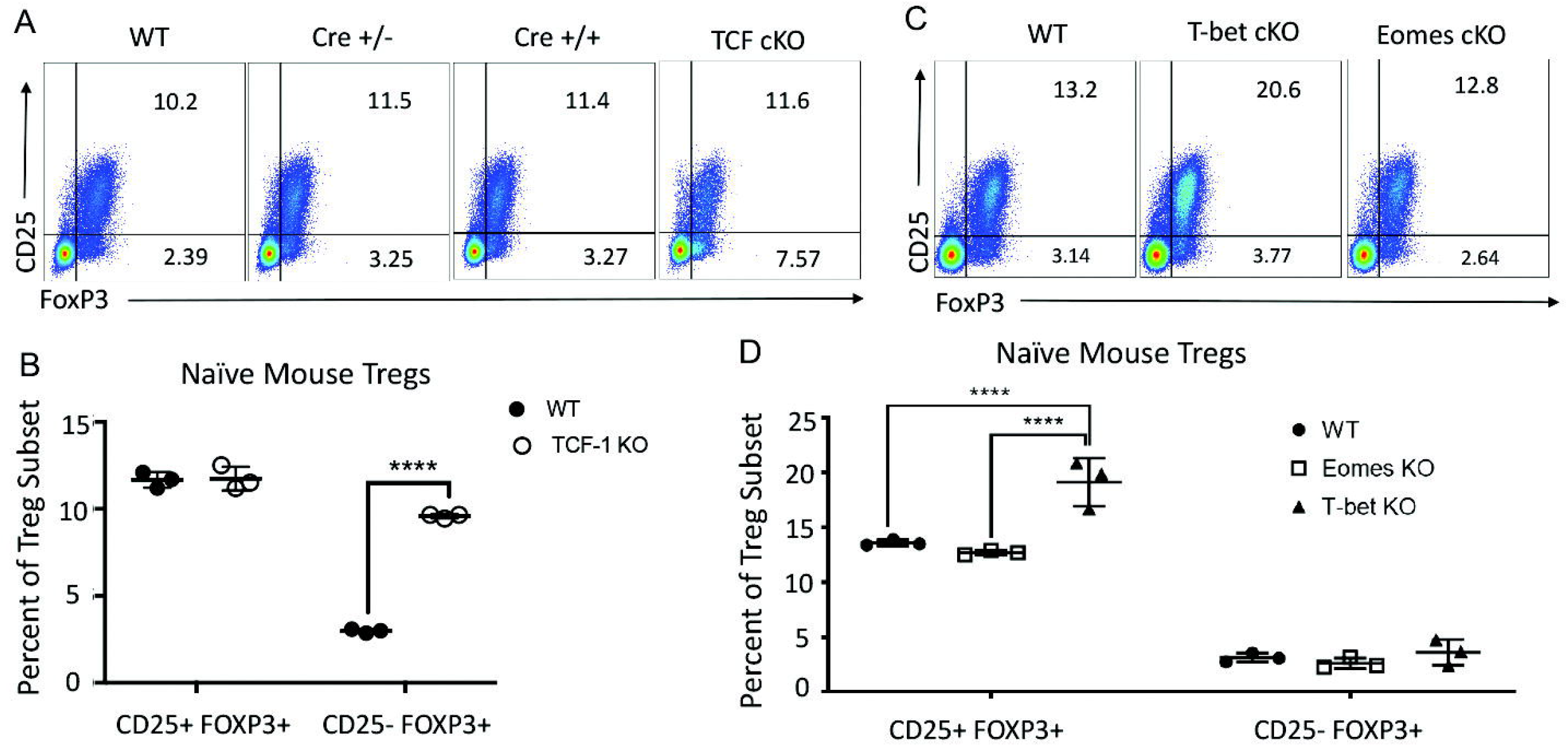
Loss of TCF-1 in T cells leads to increased production of noncanonical Tregs, with no effect by Eomes and T-bet. Splenocytes were taken from naive mice and stained for CD4, CD25, and FOXP3 to identify Treg populations. Canonical Tregs are CD4+ CD25+ FOXP3+, and noncanonical Tregs are CD4+ CD25- FOXP3+. (A) Flow cytometry plots of Treg frequencies in WT, CD4cre heterozygous control, CD4cre homozygous control, and TCF cKO naive mice, shown out of CD4 T cells. One representative plot is shown per group. (B) Quantification of (A), with CD4cre groups not shown. (C) Flow cytometry plots of Treg frequencies in WT, T-bet cKO, or Eomes cKO naive mice, shown out of CD4 T cells. One representative plot is shown per group. (D) Quantification of (C), showing all groups. For all graphs, n=3 per group and one representative experiment is shown, mean and SD are also plotted. P-values: ****=p≤0.0001, and N.S. (p>0.05) is not shown.

### Downstream factors Eomes and T-bet do not impact noncanonical Treg frequency

TCF-1 controls the T cell downstream transcription factor Eomesodermin (Eomes), and may affect the factor T-box transcription factor 21 (T-bet) through control of the T_FH_/Th1 axis^32, 40^. To examine whether changes in Eomes and T-bet during TCF-1 deficiency could play a role in the expansion of noncanonical Tregs, we performed the same phenotyping using Eomes cKO and T-bet cKO mice [Fig 1C-D]. Canonical Tregs were found to be increased in the T-bet cKO, which has been previously reported [refs], [Fig 1C-D]. However, noncanonical Tregs were not impacted by loss of either Eomes or T-bet, suggesting that these factors are not critical for expansion or suppression of these cells as TCF-1 is [Fig 1C-D]. Therefore, TCF-1 has a direct impact on the frequency of noncanonical Tregs.

### Expansion of noncanonical Tregs due to TCF-1 deficiency is cell-intrinsic

Changes in phenotype may be cell-intrinsic (due to loss of the factor in each individual cell) or cell-extrinsic (from changes in the microenvironment in the mouse due to loss of the factor). We created bone marrow chimeras to test whether the change to noncanonical Tregs was cell-intrinsic or not. Briefly, bone marrow from WT and TCF cKO mice was mixed at a 1:4 (WT:TCF cKO) ratio. The ratio was chosen based on our previous studies and observations that TCF cKO T cells do not proliferate well. The bone marrow mixture was transplanted into irradiated Thy1.1 mice (donor and host both on H2Kb background), and the mice were checked at 9 weeks by flow cytometry on blood to ensure reconstitution. At 10 weeks, splenocytes were taken from these mice and phenotyped by flow cytometry [Fig 2]. We found that TCF cKO donor T cells in the chimeras still showed increased frequency of noncanonical Tregs, despite development in a normal WT thymus [Fig 2]. Interestingly, the frequency of canonical Tregs was also increased in TCF cKO donor cells in the chimeras [Fig 2]. These results show that TCF-1 cell-intrinsically controls expansion of noncanonical Tregs.

**Fig 2.**
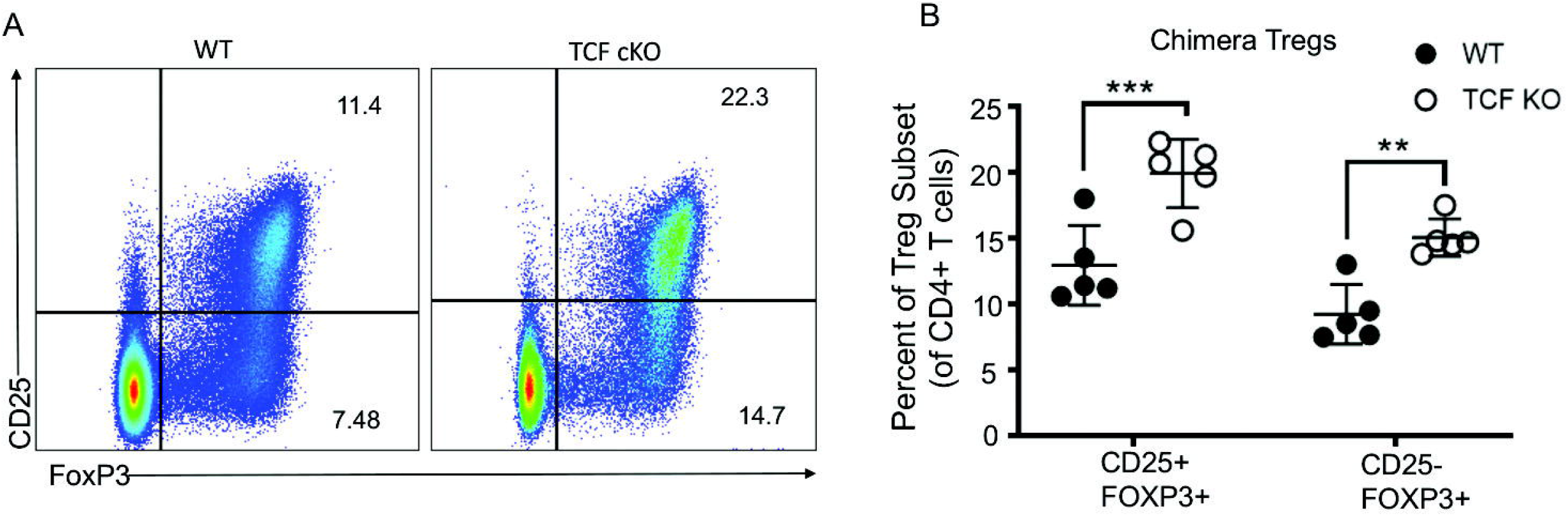
Expansion of noncanonical Tregs due to TCF-1 deficiency is cell-intrinsic. Bone marrow cells from naive WT and TCF cKO mice were obtained and mixed at a 4:1 (TCF:WT) ratio, and injected into lethally irradiated Thy1.1 mice. At 10 weeks post-transplant, flow cytometry was performed to look at Treg markers (CD25, FOXP3). (A) Flow cytometry plots of Treg frequencies from WT donor-derived or TCF cKO donor-derived CD4 T cells within the chimeric mouse. One representative plot is shown per group. (B) Quantification of (A). For all graphs, n=5 per group and one representative experiment is shown, mean and SD are also plotted. P-values: **=p≤0.01, ***=p≤0.001, and N.S. (p>0.05) is not shown.

### TCF-1-defìcient T cells induce increased production of WT noncanonical Tregs in a prionlike manner

We performed the chimera experiment described above to determine whether the effect of TCF-1 deficiency on noncanonical Tregs was cell-intrinsic. Given that the increase in CD25- Tregs was cell-intrinsic, we next sought to determine whether the number of cells was impacted along with the frequency of these cells. Loss of TCF-1 in this model results in a lower frequency and number of CD4 T cells; thus, we wanted to know if the number of noncanonical Tregs was actually higher than in WT mice, given that the frequency is based out of CD4 T cells. When we looked at the number of CD25+ FOXP3+ Tregs per 100,000 CD3 T cells in naïve WT or TCF cKO mice, we found that the number of canonical Tregs was significantly decreased in the TCF cKO mouse [Fig 3A]. This was expected given the drop in CD4 T cell numbers with no difference in canonical Treg frequency [Fig 1B]. Interestingly, the number of noncanonical (CD25- FOXP3+) Tregs was significantly higher in TCF cKO mice compared to WT mice [Fig 3A], despite the reduction in CD4 T cells. Therefore, even among total T cells, noncanonical Tregs make up a greater component of the T cells in TCF cKO mice than in WT mice. When we looked at the cell numbers per 100,000 CD4+ T cells for naive mice, we saw an increase in CD25- Treg numbers from TCF cKO mice, as expected based on the increased frequency out of CD4 T cells [Fig 3B].

**Fig 3.**
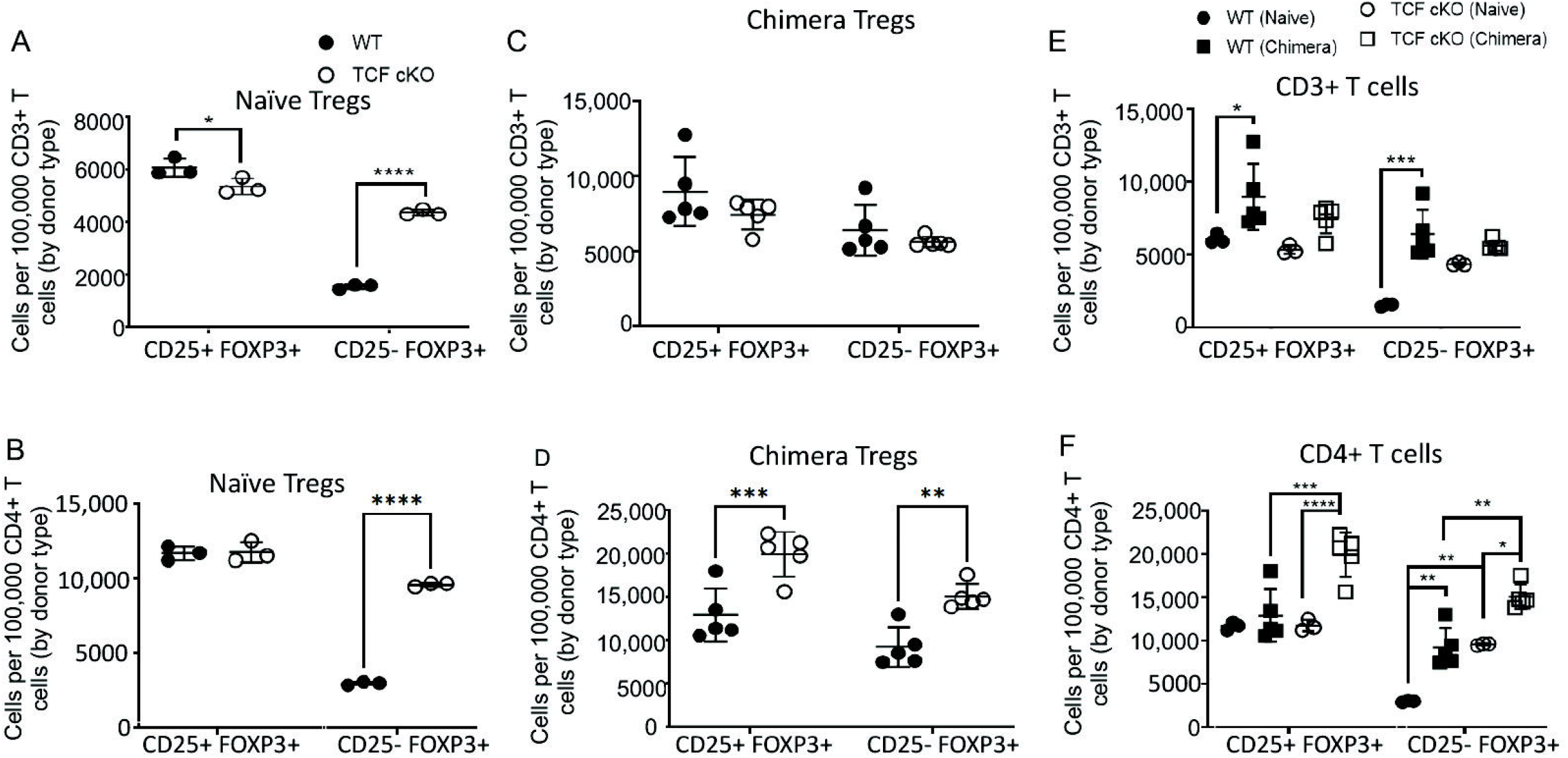
TCF-1-deficient T cells induce increased production of WT noncanonical Tregs in a prion-like manner. Splenocytes from WT or TCF cKO naive or chimeric mice were obtained and stained for flow cytometry (data shown in Fig 1 and Fig 2, respecitvely). Here, cell numbers are shown from these experiments. (A) Number of CD25+ or CD25- Tregs calculated per 100,000 CD3+ T cells for naive WT or TCF cKO mice. (B) Number of CD25+ or CD25- Tregs calculated per 100,000 CD4 T cells for naive WT or TCF cKO mice. (C) Number of CD25+ or CD25- Tregs calculated per 100,000 CD3+ WT or TCF cKO donor-derived T cells within chimeric mice. (D) Number of CD25+ or CD25- Tregs calculated per 100,000 CD4+ WT or TCF cKO donor-derived T cells within chimeric mice. (E) Data from (A) and (C) combined, showing number of CD25+ or CD25- Tregs calculated per 100,000 CD3+ T cells from WT or TCF cKO donor-derived T cells within chimeric mice, or from WT or TCF cKO naive mice. (F) Data from (B) and (D) combined, showing number of CD25+ or CD25- Tregs calculated per 100,000 CD4+ T cells from WT or TCF cKO donor-derived T cells within chimeric mice, or from WT or TCF cKO naive mice. Data quantitated from flow cytometry plots. For all graphs, n=3 to 5 per group and one representative experiment is shown, mean and SD are also plotted. P-values: *=p≤0.05, **=p≤0.01, ***=p≤0.001, ****=p≤0.0001, and N.S. (p>0.05) is not shown.

Next, we looked at the cell numbers for these Tregs in the chimeric mice. We calculated the number of Treg cells per 100,000 CD3+ T cells of each donor type (because WT and TCF cKO donor T cells were mixed in the mouse). In the chimera, we found no difference in the numbers of canonical or noncanonical Tregs in these two donor types out of CD3 T cells [Fig 3C]. However, the number of WT CD4 T cells in chimeric mice was nearly four times higher than the number of TCF cKO CD4 T cells. This could result in an increased number of Tregs seen among CD3 T cells even if the frequency of Tregs was the same for WT as for TCF cKO mice, because the number of CD4 T cells was greater per 100,000 CD3 T cells. Therefore, we looked at the number of Tregs per 100,000 CD4+ T cells, and we saw a reduced number of both CD25+ and CD25- Tregs from WT donors compared to TCF cKO donors [Fig 3D]. However, the number of noncanonical Tregs derived from WT T cells in the chimera was increased compared to the numbers found in the WT naive mice [Fig 3E,F], as was the frequency of these cells [Fig 1A, 2B]. The number of WT-derived CD25- Tregs out of CD4 T cells was increased in the chimera to the average numbers found in the TCF cKO naive mice [Fig 3F]. This is highly important as it suggests that, when in a mixed environment with WT cells, the TCF cKO T cells induce more WT T cells to adopt the noncanonical Treg phenotype, resulting in a higher frequency and number of these Tregs than would occur in naive WT mice. Therefore, loss of TCF-1 produces a prion-like ability in T cells to promote nearby cells to develop a Treg phenotype.

### Noncanonical Tregs are found at increased frequency in multiple tissues from TCF-deficient mice

The noncanonical Treg population we observed in this model was identified in the spleen. To determine whether these cells existed only in the spleen, or could be found in other organs, we phenotyped lymphocytes from the spleen, thymus, liver, and small intestine of WT or TCF cKO mice [Fig 4]. We found that noncanonical Tregs were present even in WT mice in all of the tested organs [Fig 4A]. Significantly higher frequencies of noncanonical Tregs were observed in the thymus, spleen, and liver of TCF cKO mice compared to WT mice [Fig 4B]. It was very difficult to obtain total lymphocytes from the guts of TCF cKO mice [Fig 4A]. Thus, we were unable to clearly determine whether noncanonical Tregs are present in higher frequency in the guts. Therefore, noncanonical Tregs are present in multiple tissues in WT and TCF cKO mice, with higher frequencies appearing when TCF-1 is lost. Additionally, the thymus of TCF cKO mice showed an increase in canonical Tregs [Fig 4C]. Overall, these data show that CD25- Tregs are found in multiple tissues, with expansion due to loss of TCF-1 occurring in these peripheral tissues as well.

**Fig 4.**
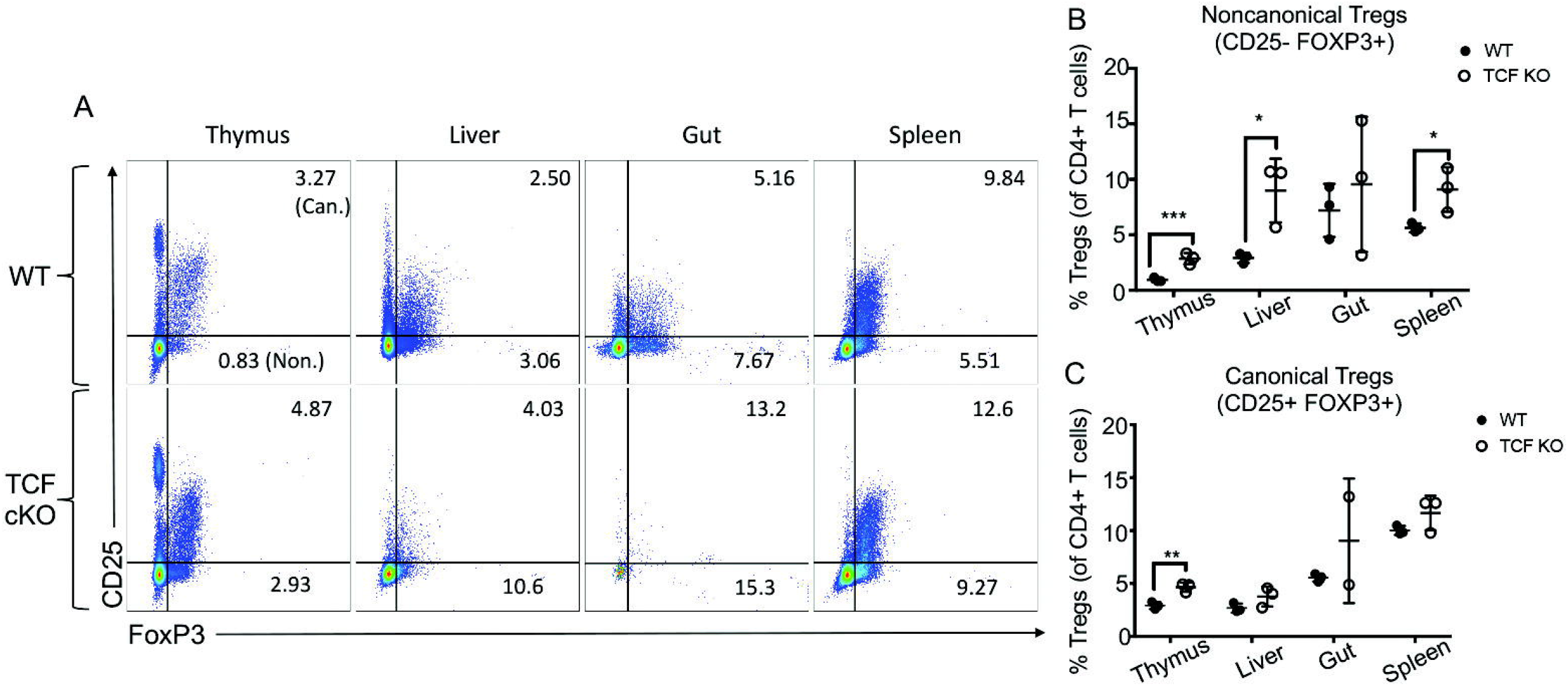
Noncanonical Tregs are found at increased frequency in multiple tissues from TCF-deficient mice. Lymphocytes were obtained from the spleen, thymus, small intestine (“gut”), and liver as described in detail in the Methods section. These cells were taken from WT or TCF cKO naive mice, and stained for flow cytometry. (A) Flow cytometry plots of Treg frequencies in tissues of WT or TCF cKO naive mice, shown out of CD4 T cells. One representative plot is shown per group. (B,C) Quantification of (A), for canonical Tregs (B) and noncanonical Tregs (C). For all graphs, n=3 per group and one representative experiment is shown, mean and SD are also plotted. P-values: *=p≤0.05, **=p≤0.01, ***=p≤0.001, and N.S. (p>0.05) is not shown.

### Expansion of noncanonical Tregs is not due to aberrant FOXP3 expression

Delacher et al. [2020]^24^ recently claimed that global deficiency of TCF-1 led to aberrant expression of FOXP3 in both CD4 and CD8 T cells, resulting in conventional T cells which appeared to be Tregs but were not. To determine whether aberrant FOXP3 expression was the reason for expansion of noncanonical Tregs, we first examined FOXP3 expression on CD8 T cells in WT and TCF cKO mice [Fig 5]. We found that there was no significant increase in FOXP3+ CD25- cells among CD8 T cells from TCF cKO naive mice [Fig 5A-B]. Therefore, aberrant expression of FOXP3 in CD8 T cells is not occurring in this model of TCF-1 deletion.

**Fig 5.**
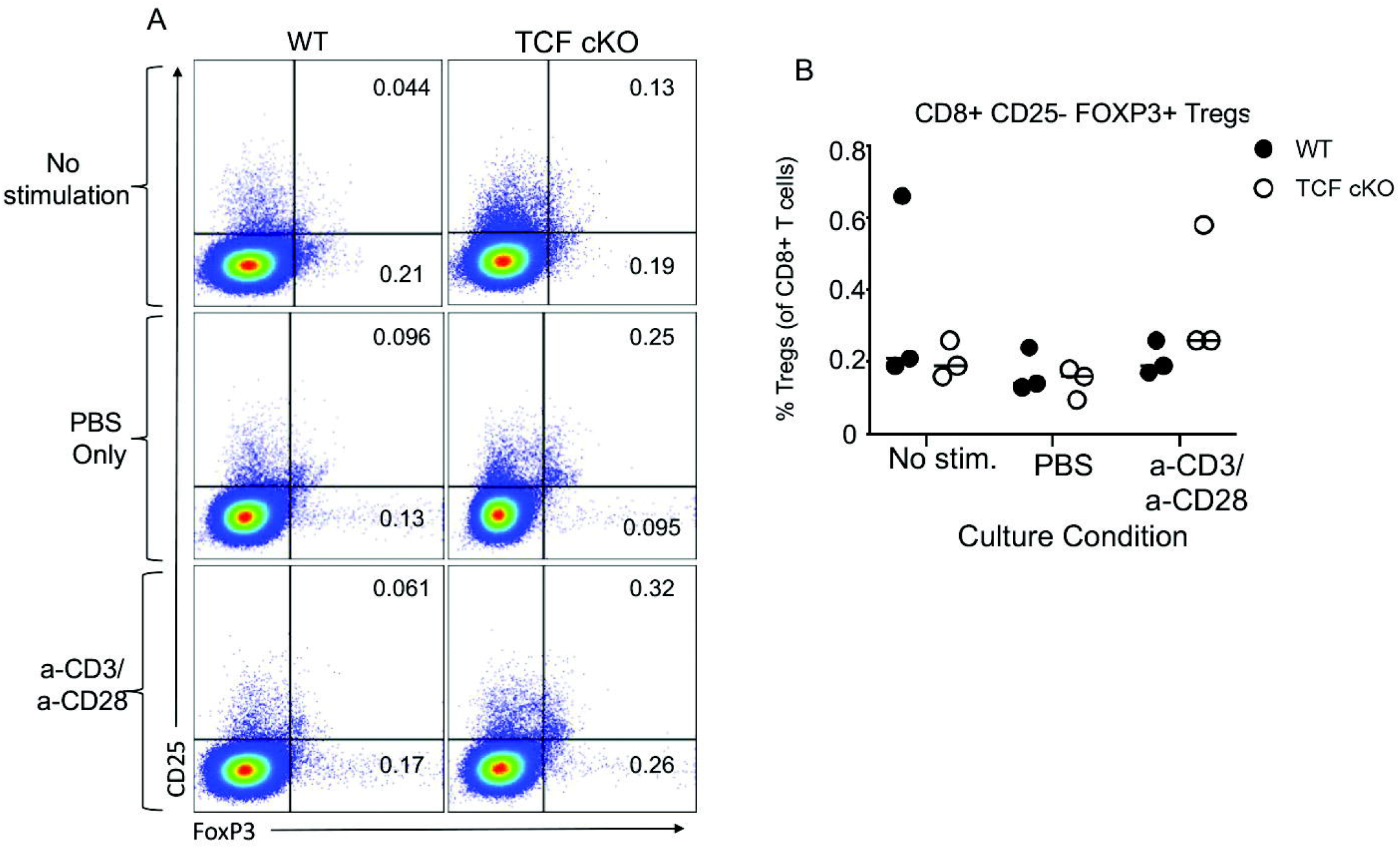
Aberrant FOXP3 expression is not visible in CD8 T cells. Splenocytes were taken from naive mice and either directly stained or cultured for 6 hours with PBS or anti-CD3/anti-CD28 stimulation prior to staining for CD4, CD8, CD25, and FOXP3. CD8 T cells were examined for expression of FOXP3 and CD25 to identify aberrant expression. (A) Flow cytometry plots of “Treg” frequencies in WT or TCF cKO naive mice, shown out of CD8 T cells. One representative plot is shown per group. (B) Quantification of (A), showing noncanonical “Treg” frequencies for each of the three conditions. For all graphs, n=3 per group and one representative experiment is shown, mean and SD are also plotted. N.S. (p>0.05) is not shown, and no significance was reported for this experiment.

Next, we examined whether CD25- Tregs from WT and TCF cKO expressed other functional Treg markers or IL-2 [Fig 6]. Briefly, splenocytes from naive mice were obtained, and split into three groups: one group was simply stained for flow cytometry, one was cultured for 6 hours with Brefeldin A in wells coated with PBS, then stained, and one was cultured for 6 hours with Brefeldin A in wells coated with anti-CD3 and soluble anti-CD28 (in the media), then stained. The uncultured samples were stained for CTLA-4, while the cultured cells were stained for CTLA-4, IL-2, and IL-10. For CTLA-4 expression, no significant difference between CD25- Tregs from WT or TCF cKO mice detected. However, Tregs from TCF cKO mice trended towards increasing frequency of CTLA-4 expression for all conditions [Fig 6A]. No significant difference was observed for IL-2 or IL-10 expression between WT and TCF cKO CD25- Tregs, but in both cases, the TCF cKO cells cultured with antibody stimulation trended toward increased expression of these cytokines compared to WT cells [Fig 6B-C]. The level of IL-2 production was similar for TCF cKO CD25- Tregs compared to WT CD25- and CD25+ Tregs, which are known to be suppressive [Fig 6B]. We did note the same trends for all three markers in CD25+ Tregs, but the frequency of cells expressing these markers tended to be higher among CD25+ than CD25- Tregs [Supp Fig 1]. This suggests that loss of TCF-1 may also enhance expression of these markers in canonical Tregs. Therefore, the CD25- Tregs observed in WT and expanded in TCF cKO mice are not simply the product of aberrant FOXP3 expression. These cells express CTLA-4 and produce IL-2 and IL-10 at similar frequencies as in WT mice, suggesting that they are truly noncanonical Tregs with suppressive function and not just activated conventional T cells.

**Fig 6.**
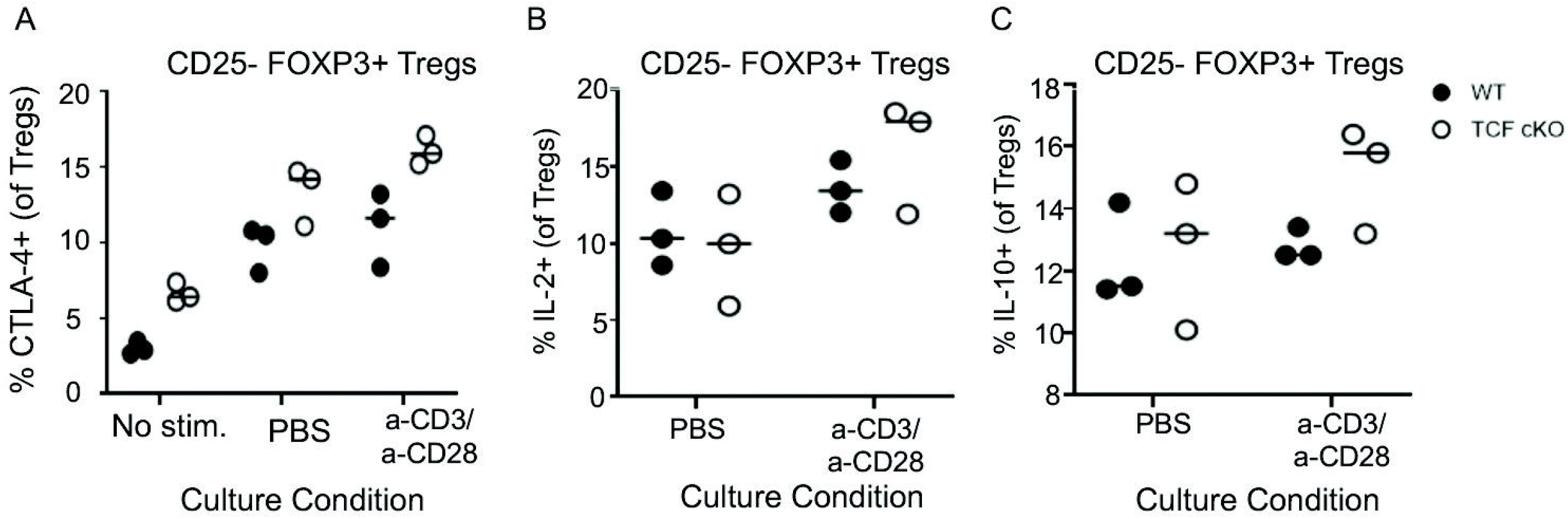
Expansion of noncanonical Tregs is not due to aberrant FOXP3 expression. Splenocytes were taken from naive mice and either directly stained or cultured for 6 hours with PBS or anti-CD3/anti-CD28 stimulation prior to staining for CTLA-4, IL-2, and IL-10. CD25- Tregs were examined for expression of these Treg markers. Flow cytometry plots not shown. (A) Quantification of CTLA-4 expression among CD25- Tregs for uncultured, control (PBS) cultured, and antibody-stimulated cultured T cells. (B) Quantification of IL-2 expression among CD25- Tregs for control (PBS) cultured and antibody-stimulated cultured T cells. (C) Quantification of IL-10 expression among CD25- Tregs for control (PBS) cultured and antibody-stimulated cultured T cells. Data quantitated from flow cytometry plots. For all graphs, n=3 per group and one representative experiment is shown, mean and SD are also plotted. N.S. (p>0.05) is not shown, and no significance was reported for this experiment.

## Discussion

TCF-1 is a major T cell transcription factor which controls T cell development and lineage, and also has an important role in T cell responses to infection^23, 24, 25, 26, 27, 28, 29, 30, 31, 32, 33, 40^. However, the role of TCF-1 in controlling mature T cells following normal development has not been well studied. Our studies sought to determine whether loss of TCF-1 in mature T cells (following development) would impact Treg cell fates in vivo in the mouse, given that TCF-1 is known to repress FOXP3 expression^24^. Loss of TCF-1 in this context – using a Tcf7^flox/flox^ x CD4cre mouse – resulted in production of a smaller number and frequency of CD4 T cells, as has been reported previously^37^. However, of the CD4 T cells present, there was a higher fraction of noncanonical Tregs found in the TCF cKO mice compared to WT, while no change in frequency was found for canonical Tregs. This suggests that the expression of FOXP3 in noncanonical cells from the TCF cKO mice may not simply be aberrant elevated expression of FOXP3, because if that were the case, one would expect more canonical Tregs as well. In addition, the expression of IL-10 and CTLA-4 at WT levels among CD25- Tregs suggests that these expanded cells from TCF cKO mice are actually potentially suppressive Treg cells, not the product of abnormal FOXP3 expression. This is in direct opposition to what Delacher et al. [2020]^24^ reported from use of global TCF-1 KO mice and CRISPR-mediated deletion of TCF-1, suggesting that TCF-1 may have a different role in mature cells than in developing cells.

Of note, our studies showed that the rise in noncanonical Tregs was cell-intrinsic, and not a result of microenvironment changes in the mouse due to loss of TCF-1, as may be more likely in a TCF-1 global KO^37^. In addition, TCF-1 controls Eomes and possibly T-bet expression^32, 40^, yet Eomes cKO and T-bet cKO mice showed no changes to noncanonical Tregs. This suggests that TCF-1 and loss thereof is directly responsible for the expansion of CD25- FOXP3+ Tregs. These noncanonical Tregs were found in small intestine, spleen, thymus, and liver, suggesting that they are capable of migrating to potential sites of immune responses where suppression may be needed. If these cells were simply aberrantly exiting the thymus without expression CD25, we would expect the frequency of these cells to be higher in the thymus than other organs, but this was not the case. In fact, the frequency of these cells was higher in liver, guts, and spleen than in thymus, especially for the TCF cKO mice. Therefore, these noncanonical Tregs appear to be stable peripheral Tregs which have lost CD25 expression at some point after exiting the thymus^13^.

Our most important observation from this study came from our mixed bone marrow chimera, which involved mixing BM cells from WT and TCF cKO mice and allowing them to develop to maturity in a WT thymus. We then examined frequency of CD25+ and CD25- Tregs in the spleen. This allowed us to determine that the expansion of CD25- Tregs is cell-intrinsic, because the rise in frequency of these cells for TCF cKO mice compared to WT mice was preserved for this model. Surprisingly, when we looked at the number of CD25+/- Tregs per 100,000 CD3 T cells of each donor type in this model, we saw no difference for either type of Treg. We thought that this could be because the number of CD4 T cells from WT donors was almost four times as high as the number of CD4 T cells from the TCF cKO donor, within each chimeric mouse. Even if the frequency of CD25- Tregs was less in WT mice, this could result in similar cell numbers when normalized to CD3 T cells. Therefore, we looked at cell numbers per 100,000 CD4 T cells. We first saw that the number of TCF cKO CD25+ Tregs in the chimera increased compared to in TCF cKO naive mice, so that the number of canonical Tregs was significantly higher than those from WT donors. This suggested that being in a WT environment induced higher production of these cells from TCF cKO cells. Unsurprisingly, we also saw that the number of CD25- Tregs was less for WT donors than for TCF cKO donors, similar to the frequency differences. However, we found that this was because the presence of TCF-1-deficient T cells had induced production of higher numbers of noncanonical Tregs from the WT donor BM than would occur in a naive WT mouse. We also saw that the frequency of these cells from WT donors was increased in the chimera compared to among naive WT T cells. This is critical as it suggests that loss of TCF-1 in T cells can alter the microenvironment (regardless of the fact that the cells in the microenvironment are WT), leading to alterations in the fate decisions and phenotype of nearby cells.

This trait is similar to the ability of pathogenic prion proteins to induce misfolding of nearby normal prion proteins into the pathogenic state^41^. In a similar manner, despite the WT environment, TCF-deficient T cells are able to induce more production of CD25- Tregs from WT donors cells. This finding – and this trait generally – could have a major impact on attempts to raise yield of suppressive cells for therapeutic purposes. It also shows a novel way by which deficiency within a cell could impact nearby cells, most likely by release of factors into the microenvironment which can alter fate decisions. To our knowledge, the ability of gene- or geneproduct-deficient cells to alter WT cells in a WT environment in this fashion has not yet been described, so this trait is a major novel finding.

Interestingly, the number of canonical Tregs was lower in TCF cKO mice when normalized to the amount of CD3 T cells, which makes sense given the drop in CD4 T cells numbers in these mice. However, despite this drop in CD4 cells, the number of noncanonical Tregs was still higher in TCF cKO mice compared to WT mice. This suggests that loss of TCF-1 increases the yield of cells adopting a suppressive fate, while potentially disrupting the expression of CD25, given that fewer CD25+ and more CD25-cells were observed. CD25 is part of the IL-2 receptor^42^, so the reduced expression of CD25 suggests that the Tregs expanded by loss of TCF-1 may not be as receptive to IL-2 stimulation, possibly providing a clue as to the apparent reduction in proliferation or survival of T cells from TCF cKO donors in the chimera model.

Overall, these results show that TCF-1 normally suppresses FOXP3 in CD4 T cells, and that loss of TCF-1-driven suppression allows expansion of CD25- FOXP3+ Tregs. These cells also express other markers found on functional Tregs, including IL-10, and CTLA-4, indicating they are true Tregs with the potential to be suppressive. The presence of TCF-deficient T cells in a WT environment induced increased production of these unique Treg from WT donor cells, showing a prion-like ability of TCF cKO T cells to control cell fate in neighboring cells in the microenvironment. These CD25- Tregs are found in multiple tissues, suggesting they may be stable Tregs rather than a product of transient aberrant FOXP3 expression. CD8 cells expressing FOXP3 but not CD25 were not increased by loss of TCF-1, further showing that these cells are not due to aberrant FOXP3 expression. In all, these data suggest that TCF-1 normally suppresses FOXP3 to control expansion of this CD25- Treg subset, and modulation of TCF-1 may provide an opportunity to increase production of these cells.

Future work should address the functional capacity of these Treg cells, although the suppressive ability of CD25- Tregs from normal WT mice has been proven in several experimental models^11, 12, 14^. The mechanism or factors by which TCF-deficient cells evoke fate changes in WT cells in a mixed cell model should also be examined, as this prion-like ability could have significant impacts on cell culture strategies for therapeutic uses.

## ABBREVIATIONS

cGys: centiGrays
cKO: conditional knock-out
Eomes: Eomesodermin
IPEX: immune dysregulation, polyendocrinopathy, and X-linked syndrome
KO: knock-out
LAK media: lymphocyte activated killer media
MHC (I/II): major histocompatibility complex (I/II)
RBCs: red blood cells
T-bet: T-box transcription factor TBX21
TCF-1/TCF: T Cell Factor-1
Tregs: regulatory T cells
WT: wild-type

## Author Contributions

RH and MK designed the experiments, analyzed the data, and wrote the manuscript. RH performed all experiments. MM assisted with experiments. MK assisted with scientific and technical design and discussion.

## Acknowledgments

We would like to thank Dr. Jyoti Misra Sen from the NIH for providing us with TCF cKO mice to start our colony. We would also like to thank members of the Karimi lab for their helpful discussions, and members of the Yang lab for proofreading this manuscript.

## Funding

This research was funded in part by a grant from the National Blood Foundation Scholar Award to (MK) and the National Institutes of Health (NIH LRP #L6 MD0010106 and AI130182 to MK.

## Conflict of interest statement

The authors have declared that no conflict of interest exists.

## Summary Figure

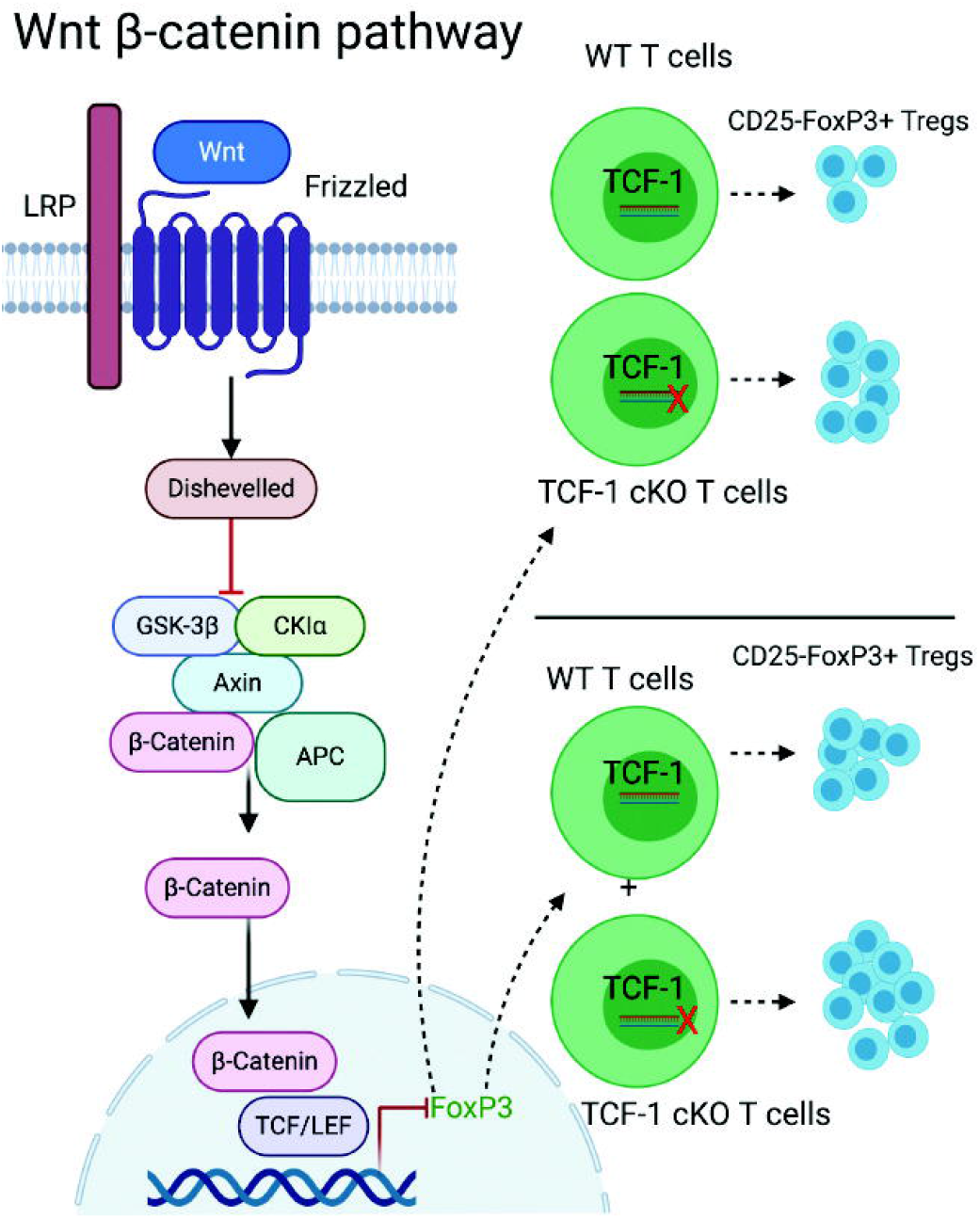

## Notes

### Competing Interest Statement

The authors have declared no competing interest.

## References

1. Li, Y. & Mariuzza, R. Structural and Biophysical Insights into the Role of CD4 and CD8 in T Cell Activation. Frontiers in Immunology 4 (2013).

2. Xing, Y. & Hogquist, K.A. T-cell tolerance: central and peripheral. Cold Spring Harb Perspect Biol 4, a006957 (2012).

3. Kappler, J.W., Roehm, N. & Marrack, P. T cell tolerance by clonal elimination in the thymus. Cell 49, 273–280 (1987).

4. Kisielow, P., Blüthmann, H., Staerz, U.D., Steinmetz, M. & von Boehmer, H. Tolerance in T-cell-receptor transgenic mice involves deletion of nonmature CD4+8+ thymocytes. Nature 333, 742–746 (1988).

5. ElTanbouly, M.A. & Noelle, R.J. Rethinking peripheral T cell tolerance: checkpoints across a T cell’s journey. Nature Reviews Immunology (2020).

6. Jørgensen, N., Persson, G. & Hviid, T.V.F. The Tolerogenic Function of Regulatory T Cells in Pregnancy and Cancer. Frontiers in Immunology 10 (2019).

7. Brunkow, M.E., Jeffery, E.W., Hjerrild, K.A., Paeper, B., Clark, L.B., Yasayko, S.A., Wilkinson, J.E., Galas, D., Ziegler, S.F. & Ramsdell, F. Disruption of a new forkhead/winged-helix protein, scurfin, results in the fatal lymphoproliferative disorder of the scurfy mouse. Nat Genet 27, 68–73 (2001).

8. Godfrey, V.L., Wilkinson, J.E. & Russell, L.B. X-linked lymphoreticular disease in the scurfy (sf) mutant mouse. Am J Pathol 138, 1379–1387 (1991).

9. Ochs, H.D., Gambineri, E. & Torgerson, T.R. IPEX, FOXP3 and regulatory T-cells: a model for autoimmunity. Immunol Res 38, 112–121 (2007).

10. Sakaguchi, S., Wing, K., Onishi, Y., Prieto-Martin, P. & Yamaguchi, T. Regulatory T cells: how do they suppress immune responses? International Immunology 21, 1105–1111 (2009).

11. Angerami, M.T., Suarez, G.V., Vecchione, M.B., Laufer, N., Ameri, D., Ben, G., Perez, H., Sued, O., Salomón, H. & Quiroga, M.F. Expansion of CD25-Negative Forkhead Box P3-Positive T Cells during HIV and Mycobacterium tuberculosis Infection. Frontiers in Immunology 8 (2017).

12. Coleman, M.M., Finlay, C.M., Moran, B., Keane, J., Dunne, P.J. & Mills, K.H. The immunoregulatory role of CD4^+^ FoxP3^+^ CD25^-^ regulatory T cells in lungs of mice infected with Bordetella pertussis. FEMS Immunol Med Microbiol 64, 413–424 (2012).

13. Curotto de Lafaille, M.A., Lino, A.C., Kutchukhidze, N. & Lafaille, J.J. CD25- T cells generate CD25+Foxp3+ regulatory T cells by peripheral expansion. J Immunol 173, 7259–7268 (2004).

14. Annacker, O., Pimenta-Araujo, R., Burlen-Defranoux, O., Barbosa, T.C., Cumano, A. & Bandeira, A. CD25- CD4+ T Cells Regulate the Expansion of Peripheral CD4 T Cells Through the Production of IL-10. The Journal of Immunology 166, 3008 (2001).

15. Apostolou, I., Sarukhan, A., Klein, L. & von Boehmer, H. Origin of regulatory T cells with known specificity for antigen. Nat Immunol 3, 756–763 (2002).

16. Furtado, G.C., Olivares-Villagómez, D., Curotto de Lafaille, M.A., Wensky, A.K., Latkowski, J.A. & Lafaille, J.J. Regulatory T cells in spontaneous autoimmune encephalomyelitis. Immunol Rev 182, 122–134 (2001).

17. Stephens, L.A. & Mason, D. CD25 Is a Marker for CD4+ Thymocytes That Prevent Autoimmune Diabetes in Rats, But Peripheral T Cells with This Function Are Found in Both CD25+ and CD25– Subpopulations. The Journal of Immunology 165, 3105–3110 (2000).

18. Dowling, M.R., Kan, A., Heinzel, S., Marchingo, J.M., Hodgkin, P.D. & Hawkins, E.D. Regulatory T Cells Suppress Effector T Cell Proliferation by Limiting Division Destiny. Front Immunol 9, 2461 (2018).

19. Jain, N., Nguyen, H., Chambers, C. & Kang, J. Dual function of CTLA-4 in regulatory T cells and conventional T cells to prevent multiorgan autoimmunity. Proceedings of the National Academy of Sciences 107, 1524 (2010).

20. Verhagen, J., Gabryšová, L., Shepard, E.R. & Wraith, D.C. CTLA-4 Modulates the Differentiation of Inducible Foxp3+ Treg Cells but IL-10 Mediates Their Function in Experimental Autoimmune Encephalomyelitis. PLOS ONE 9, e108023 (2014).

21. Zhao, H., Liao, X. & Kang, Y. Tregs: Where We Are and What Comes Next? Frontiers in Immunology 8 (2017).

22. Wan, Y.Y. & Flavell, R.A. ‘Yin-Yang’ functions of transforming growth factor-beta and T regulatory cells in immune regulation. Immunological reviews 220, 199–213 (2007).

23. Yang, B.H., Wang, K., Wan, S., Liang, Y., Yuan, X., Dong, Y., Cho, S., Xu, W., Jepsen, K., Feng, G.S., Lu, L.F., Xue, H.H. & Fu, W. TCF1 and LEF1 Control Treg Competitive Survival and Tfr Development to Prevent Autoimmune Diseases. Cell Rep 27, 3629–3645.e3626 (2019).

24. Delacher, M., Barra, M.M., Herzig, Y., Eichelbaum, K., Rafiee, M.-R., Richards, D.M., Träger, U., Hofer, A.-C., Kazakov, A., Braband, K.L., Gonzalez, M., Wöhrl, L., Schambeck, K., Imbusch, C.D., Abramson, J., Krijgsveld, J. & Feuerer, M. Quantitative Proteomics Identifies TCF1 as a Negative Regulator of Foxp3 Expression in Conventional T Cells. iScience 23, 101127–101127 (2020).

25. Chen, Z., Ji, Z., Ngiow, S.F., Manne, S., Cai, Z., Huang, A.C., Johnson, J., Staupe, R.P., Bengsch, B., Xu, C., Yu, S., Kurachi, M., Herati, R.S., Vella, L.A., Baxter, A.E., Wu, J.E., Khan, O., Beltra, J.C., Giles, J.R., Stelekati, E., McLane, L.M., Lau, C.W., Yang, X., Berger, S.L., Vahedi, G., Ji, H. & Wherry, E.J. TCF-1-Centered Transcriptional Network Drives an Effector versus Exhausted CD8 T Cell-Fate Decision. Immunity 51, 840–855.e845 (2019).

26. Johnson, J.L., Georgakilas, G., Petrovic, J., Kurachi, M., Cai, S., Harly, C., Pear, W.S., Bhandoola, A., Wherry, E.J. & Vahedi, G. Lineage-Determining Transcription Factor TCF-1 Initiates the Epigenetic Identity of T Cells. Immunity 48, 243–257.e210 (2018).

27. Kim, C., Jin, J., Weyand, C.M. & Goronzy, J.J. The Transcription Factor TCF1 in T Cell Differentiation and Aging. Int J Mol Sci 21 (2020).

28. Rutishauser, R.L., Deguit, C.D.T., Hiatt, J., Blaeschke, F., Roth, T.L., Wang, L., Raymond, K., Starke, C.E., Mudd, J.C., Chen, W., Smullin, C., Matus-Nicodemos, R., Hoh, R., Krone, M., Hecht, F.M., Pilcher, C.D., Martin, J.N., Koup, R.A., Douek, D.C., Brenchley, J.M., Sékaly, R.-P., Pillai, S.K., Marson, A., Deeks, S.G., McCune, J.M. & Hunt, P.W. TCF-1 regulates the stem-like memory potential of HIV-specific CD8+ T cells in elite controllers. bioRxiv, 2020.2001.2007.894535 (2020).

29. Wang, Y., Hu, J., Li, Y., Xiao, M., Wang, H., Tian, Q., Li, Z., Tang, J., Hu, L., Tan, Y., Zhou, X., He, R., Wu, Y., Ye, L., Yin, Z., Huang, Q. & Xu, L. The Transcription Factor TCF1 Preserves the Effector Function of Exhausted CD8 T Cells During Chronic Viral Infection. Frontiers in Immunology 10 (2019).

30. Welten, S.P.M., Yermanos, A., Baumann, N.S., Wagen, F., Oetiker, N., Sandu, I., Pedrioli, A., Oduro, J.D., Reddy, S.T., Cicin-Sain, L., Held, W. & Oxenius, A. Tcf1+ cells are required to maintain the inflationary T cell pool upon MCMV infection. Nature Communications 11, 2295 (2020).

31. Yu, Q., Sharma, A. & Sen, J.M. TCF1 and beta-catenin regulate T cell development and function. Immunol Res 47, 45–55 (2010).

32. Zhou, X., Yu, S., Zhao, D.M., Harty, J.T., Badovinac, V.P. & Xue, H.H. Differentiation and persistence of memory CD8(+) T cells depend on T cell factor 1. Immunity 33, 229–240 (2010).

33. Verbeek, S., Izon, D., Hofhuis, F., Robanus-Maandag, E., te Riele, H., van de Wetering, M., Oosterwegel, M., Wilson, A., MacDonald, H.R. & Clevers, H. An HMG-box-containing T-cell factor required for thymocyte differentiation. Nature 374, 70–74 (1995).

34. Cho, S.W., Kim, S., Kim, Y., Kweon, J., Kim, H.S., Bae, S. & Kim, J.-S. Analysis of off-target effects of CRISPR/Cas-derived RNA-guided endonucleases and nickases. Genome Res 24, 132–141 (2014).

35. Wang, Y., Wang, M., Zheng, T., Hou, Y., Zhang, P., Tang, T., Wei, J. & Du, Q. Specificity profiling of CRISPR system reveals greatly enhanced off-target gene editing. Scientific Reports 10, 2269 (2020).

36. Zhang, X.-H., Tee, L.Y., Wang, X.-G., Huang, Q.-S. & Yang, S.-H. Off-target Effects in CRISPR/Cas9-mediated Genome Engineering. Molecular Therapy - Nucleic Acids 4, e264 (2015).

37. Steinke, F.C., Yu, S., Zhou, X., He, B., Yang, W., Zhou, B., Kawamoto, H., Zhu, J., Tan, K. & Xue, H.-H. TCF-1 and LEF-1 act upstream of Th-POK to promote the CD4(+) T cell fate and interact with Runx3 to silence Cd4 in CD8(+) T cells. Nature immunology 15, 646–656 (2014).

38. Ariga, H., Shimohakamada, Y., Nakada, M., Tokunaga, T., Kikuchi, T., Kariyone, A., Tamura, T. & Takatsu, K. Instruction of naive CD4+ T-cell fate to T-bet expression and T helper 1 development: roles of T-cell receptor-mediated signals. Immunology 122, 210–221 (2007).

39. Jenner, R.G., Townsend, M.J., Jackson, I., Sun, K., Bouwman, R.D., Young, R.A., Glimcher, L.H. & Lord, G.M. The transcription factors T-bet and GATA-3 control alternative pathways of T-cell differentiation through a shared set of target genes. Proceedings of the National Academy of Sciences 106, 17876 (2009).

40. Shao, P., Li, F., Wang, J., Chen, X., Liu, C. & Xue, H.-H. Cutting Edge: Tcf1 Instructs T Follicular Helper Cell Differentiation by Repressing Blimp1 in Response to Acute Viral Infection. The Journal of Immunology 203, 801 (2019).

41. Kupfer, L., Hinrichs, W. & Groschup, M.H. Prion protein misfolding. Curr Mol Med 9, 826–835 (2009).

42. Létourneau, S., Krieg, C., Pantaleo, G. & Boyman, O. IL-2- and CD25-dependent immunoregulatory mechanisms in the homeostasis of T-cell subsets. J Allergy Clin Immunol 123, 758–762 (2009).

